# Pain and touch differentially modulate corticospinal excitability, independent of afferent inhibition

**DOI:** 10.1101/2025.05.09.653032

**Authors:** Louisa Gwynne, Luigi Tamè

## Abstract

Pain can profoundly impact motor functioning to support self-preservation but is also associated with motor and somatosensory disturbances. Despite considerable research exploring the influence of pain and touch on motor and sensorimotor processes, the nature of this relationship remains elusive. Specifically, it is uncertain whether pain and touch modulate motor processes independently of each other or are interconnected. Across two experiments, an afferent inhibition (AI) paradigm was tested to probe the effects of tactile and nociceptive inputs on corticospinal processes and sensorimotor interactions. In Experiment 1 (*N*=20), the effect of electrocutaneous stimulation duration (0.2 vs. 0.4 ms) on transcranial magnetic stimulation (TMS)-induced corticospinal excitability (CSE) was assessed using a short and long-latency AI paradigm. A single electrocutanous stimulus was delivered to the left index finger before single pulse-TMS over the right-first dorsal interosseous (FDI) motor hotspot at one of five delays (15, 25, 35, 45, 60 or 160 ms). In Experiment 2 (*N*=20), the same paradigm was used to examine if this effect of sensorimotor interaction is changed when moderate tonic heat pain is delivered to the forearm. Significant AI was observed in both experiments at delays of 25, 35 and 160 ms, with afferent facilitation at 60 ms. This effect was not influenced by the duration of afferent stimulation (Experiment 1) nor by the presence of heat pain (Experiment 2). However, we found a significant reduction in CSE in painful compared to painless conditions, indicating that while tonic pain modulates CSE, tactile afferent inhibition remains unaffected. This supports the notion that pain has a direct (inhibitory) effect on motor output; however, in this context, tactile sensorimotor interactions remain unaltered.

## Introduction

Dynamic interaction between the somatosensory and motor systems permits accurate engagement with the environment, supporting adaptive responses and object manipulation (Johansson & Flanagan, 2009; Zafarana et al., 2024). It is well-documented that somatosensory signalling can alter the functioning of the motor system, including nociceptive processing that is associated with disrupted corticospinal motor networks (Rohel et al., 2021) and contributes to sensorimotor disturbances (Harris, 1999; McCabe et al., 2005). For example, hypertonic saline-induced muscle pain alters cortical motor circuits, reducing intracortical facilitation during pain and increasing inhibition post-pain, as shown by changes in the transcranial magnetic stimulation (TMS)-evoked corticospinal excitability (CSE) (Schabrun & Hodges, 2012). However, the relationship between cortical motor and somatosensory processes in pain remains unclear.

The modulatory effects of pain on corticospinal inhibitory and excitatory networks are well explored. Experimentally induced cutaneous thermal and muscle pain are largely associated with CSE inhibition (for review see Chowdhury et al., 2022; Rohel et al., 2021). The inhibition of TMS-induced CSE has been found with both momentary (phasic) and continuous (tonic) cutaneous thermal pain (Billot et al., 2018; Cheong et al., 2003; Dubé & Mercier, 2011; Farina et al., 2001; Mavromatis et al., 2016) as well as tonic muscle pain (Burns et al., 2016a, 2016b; Larsen et al., 2018, 2019; Rice et al., 2021; Schabrun & Hodges, 2012). However, experimentally induced nociceptive pain does not produce consistent inhibitory effects on CSE (Rohel et al., 2021). For instance, neither hypertonic saline nor topical capsaicin-induced cheek pain affected CSE (Romaniello et al., 2000). Similarly, hypertonic saline-induced hand pain reduced motor excitability, whereas forearm-induced pain had no effect (Larsen et al., 2018). Furthermore, acute CO2 laser pain delivered to the hand, but not the forearm, significantly reduced CSE in the biceps brachii (Valeriani et al., 2001). Additionally, while one study found that topical capsaicin-induced tonic heat pain of the hand dorsum did not affect CSE of the ipsilateral FDI muscle (Mavromatis et al., 2016), another study reported the effect of forearm pain on motor excitability was variable from one participant to the other (Martel et al., 2017).

The motor and somatosensory cortices share reciprocal communication, with projections from the primary somatosensory cortex (S1) to the primary motor cortex (M1), predominantly inhibitory (Widener & Cheney, 1997). Insights into this intracortical connectivity have been elaborated on by the reported effects of tactile afferent stimulation on TMS-induced motor-evoked potentials (MEPs; Chen et al., 1999; Tamè & Holmes, 2023; Tokimura et al., 2000). Notably, a single preceding electrocutaneous stimulus can inhibit CSE (Bailey et al., 2016; Chen et al., 1999; Tamburin et al., 2005; Tokimura et al., 2000; Turco et al., 2017). This phenomenon depends on the temporal delay between peripheral afferent input and TMS delivery over contralateral M1, with inhibition seen at short (<40 ms) or long latencies (≥80 ms), termed short- and long-latency afferent inhibition (Chen et al., 1999; Tokimura et al., 2000). The magnitude of afferent inhibition is also influenced by the extent of sensory afferent recruitment, with increased inhibition by greater stimulation intensities (Bailey et al., 2016; Turco et al., 2017).

Nociceptive processing can further influence afferent inhibition, for example, SAI and LAI in the right first dorsal interosseous (FDI) muscle were reduced after the cessation of, but not during, saline-induced muscle pain (Burns et al., 2016b). Conversely, a more recent study found that tonic hand pain induced by cold water immersion led to reduced SAI both during immersion and immediately after limb removal and reported no influence of pain on LAI (Delahunty et al., 2024). Beyond tonic pain models, one study found that acute cutaneous heat pain had no effect on SAI when transient painful stimulation was applied to the tested hand simultaneously with the electrotactile afferent stimulus (Mercier et al., 2016).

Overall, the modulatory effects of nociceptive pain on such sensorimotor interactions remain elusive, lacking a sufficient number of investigations to attain a comprehensive understanding. Nonetheless, in this context, SAI can provide an interesting model for understanding how pain influences fundamental networks of sensorimotor processing and the nervous system’s capability to shape adaptive motor responses to threatening stimuli and physical injury. These insights can have a direct clinical relevance for understanding atypical sensory and motor functioning in pain-related disorders. Notably, SAI has been proposed as a researcher tool to understand and, even as a potential biomarker for, neurological pathologies including Parkinsons and Alzheimer’s disease (Choudhury et al., 2023; Di Lazzaro et al., 2002; Sailer et al., 2003; Yarnall et al., 2013).

To probe the effects of nociceptive pain on corticospinal processes we conducted two experiments. The first optimised a SAI paradigm to investigate the influence of somatosensory afferent stimulation delivered to the left index finger on CSE and confirm a timeline of this effect. The second experiment used the SAI paradigm to investigate the modulatory effects of tonic cutaneous thermal heat pain on corticospinal and sensorimotor functioning. Given that nociceptive-motor interactions vary by pain characteristics (Farina et al., 2003) and, the inhibitory effect on CSE by forearm pain are inconsistent (Rohel et al., 2021; Valeriani et al., 2001), the SAI paradigm was paired with a tonic model of forearm pain. This was achieved using a contact heat thermode applied on the forearm ipsilateral to the afferent stimulus (i.e., the left forearm). Based on previous reports, painful input may influence CSE through a direct pathway to the motor cortex (Chowdhury et al., 2022). In this scenario, we would expect pain to modulate CSE without affecting SAI, relative to painless conditions (Figure 1A). Alternatively, pain may exert an indirect effect on motor responses via the modulation of touch. In this case, we would predict changes in SAI, while CSE remains unaltered across the painful and painless conditions (Figure 1B). Lastly, pain may impact motor responses through both direct and indirect pathways - via the motor cortex and through touch – given that these mechanisms are not mutually exclusive (Delahunty et al., 2024). In this case, both CSE and SAI would differ between painful and painless conditions (Figure 1C).

**Figure 1.**
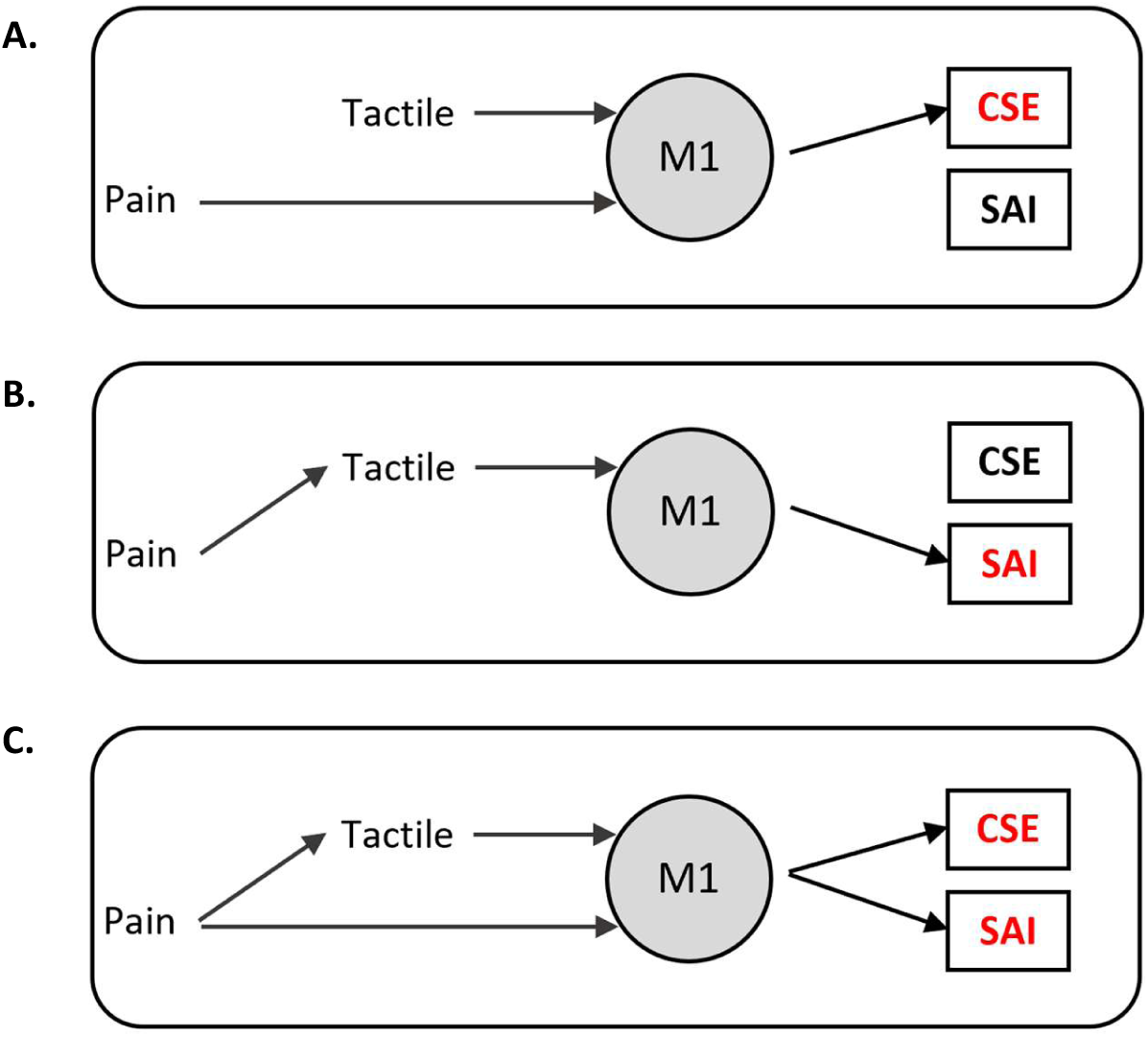
Graphical representation of proposed pathways by which ongoing pain modulates motor response during concurrent transient tactile input. (A) pain exclusively influences M1 activity altering CSE without affecting short-latency afferent inhibition (SAI). (B) Pain modulates motor response indirectly via tactile input, affecting SAI but not overall CSE. (C) Pain influences motor response through both direct (M1) and indirect (tactile-mediated) pathways, modulating both CSE and SAI.

### Experiment 1: Corticospinal excitability as a function of time and afferent duration

Experiment 1 was designed to investigate the effect of tactile input on CSE using a classical afferent inhibition paradigm (Chen et al., 1999; Tamburin et al., 2001, 2005; Tokimura et al., 2000), including any modulation of this by differing tactile durations. We hypothesised afferent inhibition at short (∼25-35 ms) and long (>100ms) latencies between stimulation of the digital nerves at the fingers and TMS-induced CSE, consistent with the current literature (Turco et al., 2018). Furthermore, we predicted that a longer afferent duration (0.2 vs 0.4 ms) would enhance SAI. This prediction was made given that longer stimulation progressively recruits additional sensory afferent fibres, while the transient nature of afferent stimulation prevents sustained changes in CSE (Chipchase et al., 2011).

## Materials and Methods

### Participants

Twenty participants were recruited *(Mean_Age_*= 20.5 *SD_Age_*= 5.6 years; 6 males, 14 females). All participants were right-handed as indicated by self-report on the Edinburgh Handedness Inventory questionnaire (EHI; Oldfield, 1971; *M* = 85, *range* = 33-67). Participants were recruited voluntarily in exchange for monetary payment at a rate of £7.50 an hour or through a university participation scheme for course credit. All participants had normal sensation to the hands and arms as indicated through free self report. Moreover, participants had no neurological, pain-related or psychiatric conditions, no current use of psychiatric and neuroactive medications or consumption of analgesic substances at least 24 hours before participation and no current use of any other prescribed medication indicated by self-report using a departmental TMS laboratory questionnaire. All procedures across experiments were approved by the School of Psychology, University of Kent ethics committee and adhered to published TMS safety guidelines (Rossi et al., 2021). A priori power analysis in G*Power (version 3.1.9.7; Faul et al., 2007) indicated that a sample of 20 participants would provide power (80%) to detect a minimum effect size (ηp2) of .060 (α = .05) for the within-subjects interaction (Delay x Stimulus Length).

### TMS and Neuronavigation

TMS procedures are reported in line with guidelines recommended for studies investigating the motor system (Chipchase et al., 2012). Biphasic single-pulse TMS was delivered with a MAG and More PowerMAG 100 Repetitive Magnetic Stimulator via a 70-mm diameter figure-of-eight coil. Electromyography (EMG) was recorded with two Ag-AgCI surface electrodes placed over the first dorsal interosseus (FDI) muscle of the left hand in a belly-tendon montage with ∼2cm inter-electrode distance. The ground electrode was placed 1-2 cm proximal to the left pisiform bone of the left wrist and skin preparation was completed using an alcohol swab. EMG signal was sampled in BrainSight Neuronavigation software (version 2.4.10; Rogue Research Inc., Montreal, QC) with a 3000 Hz sampling frequency. Scalp localisation and coil orientation were optimised using the same BrainSight Neuronavigation software based on functional MEPs and determined MNI targets. The total number of TMS pulses delivered to each participant was 280 plus an additional 50-150 pulses during the M1 localisation and TMS threshold procedures.

### M1 localisation and TMS threshold intensity estimation

TMS intensity for the main experimental protocol was set at 110% of the optimal scalp coil positioning and TMS intensity to elicit at least a 50-mV peak-to-peak motor evoked potential (MEP) in the left FDI muscle whilst at rest (resting motor threshold; RMT) in five out of 10 trials. A 3x3 grid with 10 mm spacing was drawn onto the participant’s reconfigured digitalised scalp in BrainSight, centred on our lab average right M1-FDI (at rest) coordinates from previous experiments or, in the case of Experiment 1 Session 2, the grid was centred on the participant’s right M1-FDI MNI co-ordinates determined in Experiment 1 Session 1. Starting at the central grid point, TMS was applied at 35% of the maximum stimulator output, increasing in increments of 5% until muscle evoked activity was visible on the EMG recording. The coil was moved across the grid, stimulating each grid point twice to find the optimal location (i.e. maximal MEP output); the intensity was adjusted in increments of 5% as appropriate. Next, a new 3x3 grid with 5 mm spacing was centred on this optimal location and the procedure was repeated. Finally, coil orientation was adjusted until desired MEPs were recorded. On average, the coil handle remained 45 degrees relative to the sagittal plane. Across all experiments, the mean average right M1-FDI RMT was 43.1 % (*SD* = 5.53; Table 1) and the mean average right M1-FDI MNI coordinates were 35.0, -5.1, 58.8 (x,y,z).

**Table 1.** Sensory detection thresholds (SDT) and TMS thresholds (%) across experiments and tactile conditions.

| Participant | Experiment 1 |  |  |  | Experiment 2 |  |
| --- | --- | --- | --- | --- | --- | --- |
|  | 0.4 ms |  | 0.2 ms |  | 0.2 ms |  |
|  | SDT (mA) | TMS (a.u.) | SDT (mA) | TMS (a.u.) | SDT (mA) | TMS (a.u.) |
| 1 | 1.37 | 44.0 | 1.69 | 44.5 | 1.86 | 38.0 |
| 2 | 0.77 | 39.0 | 1.17 | 41.0 | 1.25 | 36.5 |
| 3 | 0.87 | 39.5 | 1.69 | 40.0 | 0.92 | 38.5 |
| 4 | 0.95 | 43.0 | 1.73 | 45.5 | 1.42 | 51.5 |
| 5 | 0.57 | 43.5 | 0.85 | 43.0 | 1.10 | 43.0 |
| 6 | 0.73 | 50.5 | 1.01 | 51.5 | 0.97 | 50.0 |
| 7 | 0.84 | 35.5 | 1.39 | 35.5 | 0.89 | 41.0 |
| 8 | 1.29 | 43.0 | 1.77 | 41.5 | 1.44 | 57.0 |
| 9 | 1.07 | 47.5 | 1.93 | 48.0 | 1.71 | 40.0 |
| 10 | 1.15 | 38.5 | 1.81 | 39.0 | 1.23 | 52.0 |
| 11 | 0.99 | 39.5 | 1.69 | 40.0 | 1.89 | 42.5 |
| 12 | 1.02 | 45.0 | 1.64 | 47.0 | 1.61 | 49.0 |
| 13 | 1.09 | 40.5 | 1.33 | 40.5 | 2.04 | 44.5 |
| 14 | 0.84 | 43.0 | 1.45 | 39.5 | 1.11 | 49.0 |
| 15 | 1.20 | 40.5 | 2.03 | 41.0 | 1.51 | 44.5 |
| 16 | 0.73 | 48.0 | 1.34 | 46.5 | 1.20 | 29.0 |
| 17 | 0.97 | 41.0 | 1.15 | 40.5 | 2.68 | 51.0 |
| 18 | 0.88 | 40.0 | 1.41 | 40.0 | 1.33 | 37.5 |
| 19 | 1.29 | 34.5 | 1.77 | 34.5 | 0.89 | 47.5 |
| 20 | 1.11 | 48.5 | 1.65 | 48.5 | 1.29 | 34.0 |
| **Mean** | 0.99 | 42.2 | 1.52 | 42.3 | 1.42 | 43.8 |
| **SD** | 0.21 | 4.22 | 0.31 | 4.37 | 0.45 | 7.09 |

*Note. TMS thresholds are at 100% RMT.*

Innocuous transcutaneous electrical stimulation was delivered by a bipolar constant current stimulator (DS5; Digitimer, Welwyn Gardne City, United Kingdom) controlled in MATLAB (version 2019b) through a National Instruments data-acquisition card (6001). Two stainless steel electrode rings were fitted at the proximal and intermediate phalange of the participants’ left index finger. This method generated a singular, fast onset-offset tap-like sensation to the index finger by momentary activation of predominantly Aβ mechanoreceptive fibres. Square-wave pulse widths were either 0.4 ms or 0.2 ms; session 1 and 2, respectively). The participants’ electrotactile sensory detection threshold (SDT; the minimum stimulator output detectable by the participant) was estimated using a 2-down/1-up automated staircase procedure. This converges to an approximate 71% perceivability threshold (Levitt, 1971). Starting at 0.1 mA, the stimulation increased by 0.3 mA step-sizes, reduced to 0.1 mA after the first reversal and to 0.02 mA after the second reversal. The staircase ended at the eighth reversal and the SDT was calculated as the average of the last two reversals. During the main afferent inhibition experimental protocol, electrotactile stimulation was delivered at 2.5 times the SDT; all participants reported this as perceivable but not salient or painful. Table 1 displays the SDTs across experiments. For both the 0.2 ms and 0.4 ms tactile stimulation conditions, the experimental stimulation level was always calculated relative to the specific SDT of that pulse duration. As previous studies have shown that higher intensity of afferent stimulation can induce greater SAI (Bailey et al., 2016), our protocol aimed to vary the afferent timing while minimizing differences in intensity. This is reflected in the fact that the SDTs (mA) were significantly lower for the 0.4 ms condition compared to the 0.2 ms condition (*t* = 11.67, *p*<.001; Table 1).

### Experiment Protocol

M1 localisation and TMS intensity thresholding were completed first, followed by electrotactile thresholding. Lastly, the main experimental protocol was completed. Participants sat comfortably at a desk, resting their left arm and forearm in front of them in a natural position aligned with the body. The right hand and forearm were rested on the right thigh palm-down. Neither hands were covered from site throughout the experiment. Participants were instructed to fix their gaze on a black fixation cross that remained on a grey screen throughout the experiment, positioned centrally on a monitor approximately 60 cm from the participant. The experimental protocol consisted of 8 blocks, each including 35 trials and a 5-second inter-trial interval (ITI). On every trial, a single electrotactile stimulus was delivered to the left index finger at a randomly jittered timepoint between 0-1 second from the onset of the trial. Following the afferent stimulus onset, spTMS was delivered over the right M1-FDI hotspot (Figure 2). The temporal delay between afferent onset and spTMS was one of six time points (15, 25, 35, 45, 60 or 160 ms). Early delays (<100ms) were selected given the suggestion that S1 tactile-induced activity persists for at least 60 ms after afferent offset (Allison et al., 1992; Mauguière et al., 1997). Furthermore, its signal recovery time as measured by somatosensory evoked potentials (SEPs) using magnetoencephalography (MEG) has been reported at ∼110 ms (Hamada et al., 2002) and late delays are associated with long-latency AI therefore, we included a late 160ms delay. Control trials were included whereby a 0 mA electrotactile stimulus was “delivered” (TMS-only trials) followed by spTMS always at a 40 ms delay. Each block contained five trials for each of the interstimulus delays plus five control trials. The order of trial type was randomised within each block. Blocks lasted ∼4 minutes each, with 2-3 minutes breaks between them.

**Figure 2.**
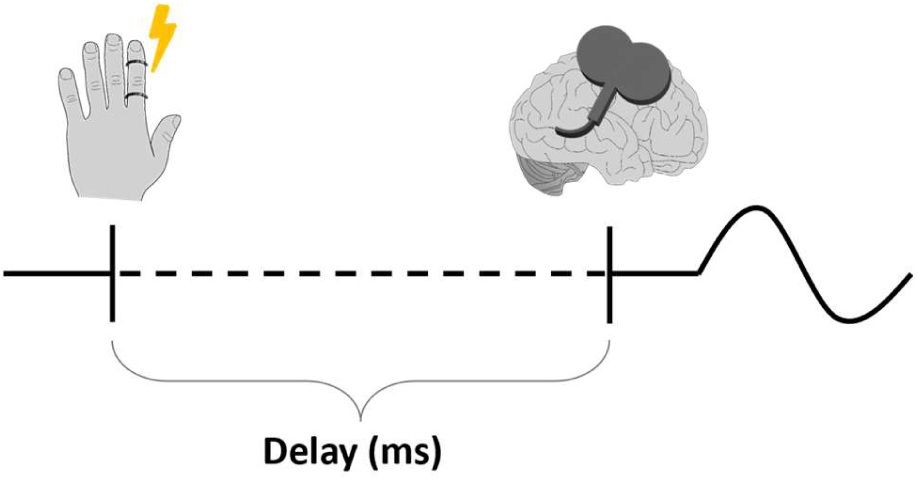
Schematic depiction of the experiment trials. An electrocutaneous stimulus was delivered to the left index finger through two ring electrodes. After afferent stimulation onset, spTMS over right-M1 was delivered at a single delay of 15, 25, 35, 45, 60 or 160 ms following a random order.

### Design

Sessions 1 and 2 were completed under a within-participants design. To assess the effect of afferent duration on SAI the factor of Delay was examined on five levels (DELAY: 15, 25, 35, 45 and 60 ms) and, the factor of afferent duration was examined on two levels (STIMULUS LENGTH: 0.4ms - session 1 vs 0.2 ms - session 2). We further investigated the overall effect of touch on CSE as a function of afferent duration (STIMULUS LENGTH: 0.4 vs. 0.2 ms) across all conditions of afferent stimulation (CONDITION; TMS-only, 15, 25, 35, 45, 60 and 160 ms delays).

## Results

MEP amplitudes were calculated as the peak-to-peak amplitude in the 10-90 ms time window after TMS pulse was delivered. MEP amplitudes at each delay were then averaged across blocks, and a MEP ratio was calculated by normalising the average MEP amplitude to the average MEP amplitude of TMS-only trials. To improve data distribution, MEP ratios were then log-transformed meaning that a MEP ratio of 0 (*SD* = 0) represented TMS-only trials. First, to test for AI we compared the MEP ratio at each delay to a MEP ratio of 0 (TMS-only). Two-tailed t-tests revealed MEP ratios were significantly less than 0 for both 0.2 and 0.4ms afferent durations at delays of 25 ms (0.2: *M* = -0.17, *SE* = 0.04; *t*(19) = 4.75, *p* = <.001, *d* = 1.06; 0.4: *M* = -0.24, *SE* = 0.03; *t*(19) = 7.15, *p* = <.001, *d* = 1.60), 35 ms (0.2: *M* = -0.09, *SE* = 0.03; *t*(19) = 3.26, *p* = .004, *d* = 0.73; 0.4: *M* = -0.18, *SE* = 0.04; *t*(19) = 4.33, *p* <.001, *d* = 0.97) and 160 ms (0.2: *M* = -0.35, *SE* = 0.05; *t*(19) = 7.00, *p* = <.001, *d* = 1.57; 0.4: *M* = -0.37, *SE* = 0.04; *t*(19) = 9.24, *p* <.001, *d* = 2.07). Differently, at a 60 ms delay, MEP ratios were significantly greater than 0 with a 0.4 ms afferent duration but not with a 0.2 ms duration (0.2: *M* = 0.09, *SE* = 0.04; *t*(19) = -2.04, *p* = .056, *d* = -0.46; 0.4: *M* = 0.13, *SE* = 0.05; *t*(19) = -2.68, *p* = .015, *d* = -0.6), indicating afferent facilitation.

A 2-way RM ANOVA (DELAY x STIMULUS LENGTH) was conducted to assess the effect of delay and stimulus duration on SAI. Because a delay of ≥100 ms is related to long-latency afferent inhibition (LAI) this timepoint was analysed separately to delays related to SAI. Mauchly’s test of sphericity indicated significantly violated compound symmetry for the factor of DELAY (*p* <.001) and a Greenhouse Geiser correction was applied. A main effect of DELAY was observed; *F*(2.13, 40.46) = 26.24, *p* <.001, η_p_^2^ = .580, reflecting both afferent inhibition and facilitation (Figure 3A). There was no significant main effect of STIMULUS LENGTH; *F*(1,19) = 1.22, *p* = .283, η_p_^2^ = .06. However, there was a significant interaction between DELAY and STIMULUS LENGTH; *F*(4, 76) = 3.00, *p* = .023, η_p_^2^ = .136. Despite this interaction, no meaningful significant differences were observed between afferent durations as a function of delay. Such interaction may have been caused by a reversal in the pattern of effects at 25 and 60 ms delays between 0.2 and 0.4 ms afferent durations (Figure 3A).

**Figure 3.**
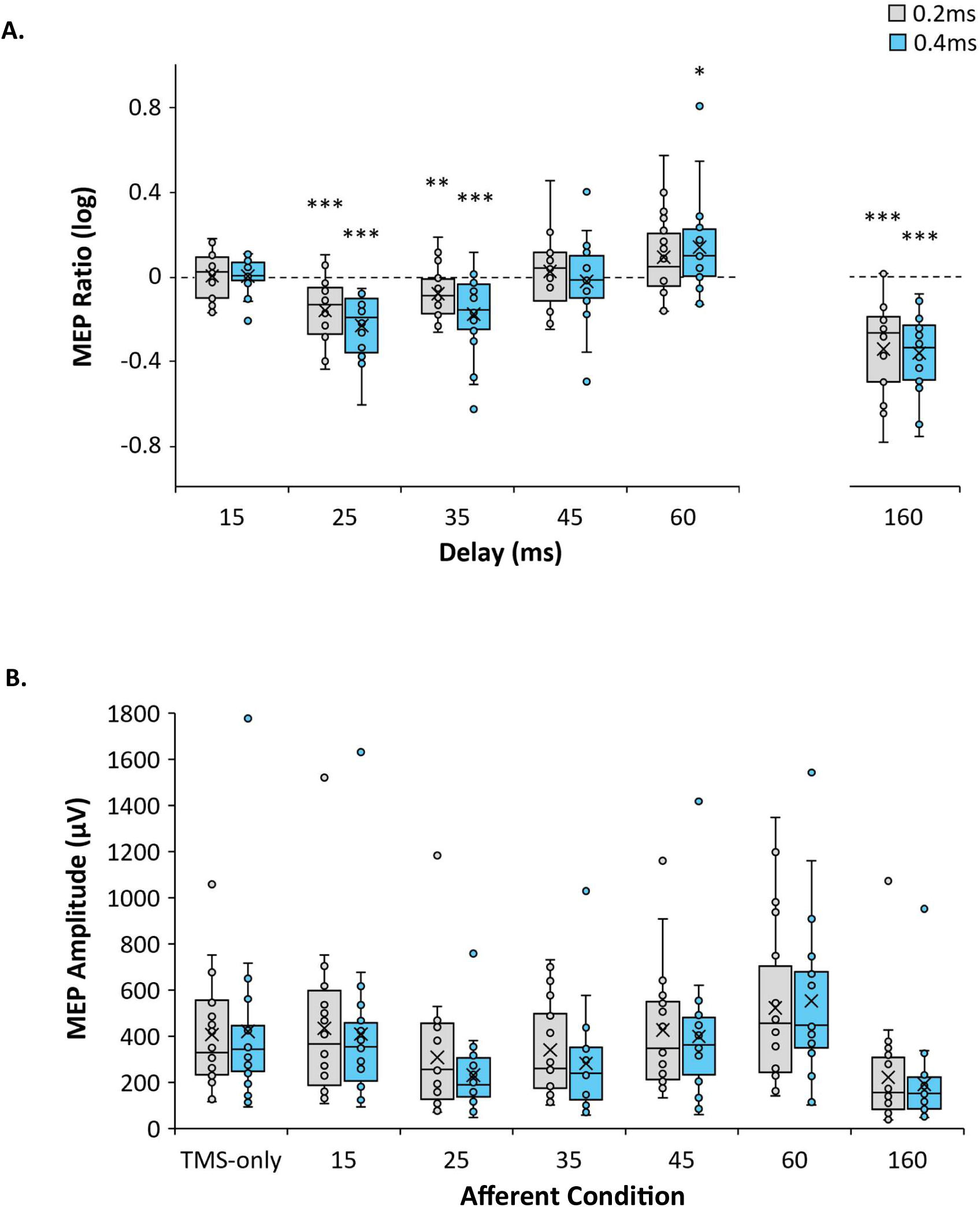
Boxplots show group data, centre crosses show the mean average and horizontal lines show exclusive median. Upper and lower whiskers show values within 1.5 times the interquartile range from lower and upper quartiles. Inner circle outlines show data points lying between inner and upper whiskers. (A) Showing MEP Ratios as a function of afferent stimulation duration and the delay (ms) between tactile input and M1-spTMS. TMS-only trials correspond to a MEP ratio of 0 and is represented by the dotted line with values above and below indicating facilitation and inhibition, respectively. \**P* < .05, \*\**P* < .01, \*\*\**P* < .001 indicate delays significantly different from a MEP ratio of 0. (B) MEP amplitudes across all trial types of afferent stimulation and TMS-only trials as a function of afferent duration.

To assess any effect of afferent duration on LAI, a two-tailed t-test was conducted to compare MEP ratios at a delay of 160 ms. This revealed no significant difference in MEP ratios between the 0.2 ms (*M* = -0.35, *SE* = 0.05) and 0.4 ms (*M* = -0.37, *SE* = 0.04) afferent durations at a 160 ms delay; *t*(19) = 0.33, *p* = .747, *d* = 0.07. For completeness, the 2-way RM ANOVA was repeated including the 160 ms delay, though, results were unaltered.

Lastly, the effect of touch on CSE was examined using a second 2-way RM ANOVA on the factors CONDITION and STIMULUS LENGTH. Mauchly’s test of sphericity indicated significantly violated compound symmetry for the factor of CONDITION (*p* <.001), so a Greenhouse Geiser correction was applied as appropriate. The analysis revealed a main effect of CONDITION; *F*(2.49, 47.27) = 21.15, *p* <.001, η_p_^2^ = .527, in that the effect of afferent inhibition and facilitation were still evident in the MEP amplitudes before ratio corrections and transformations were applied (Figure 3B). However, there was no significant main effect of STIMULUS LENGTH; *F*(1,19) = 0.29, *p* = .598, η_p_^2^ = .015, and this did not vary as a function of CONDITION (CONDITION X STIMULUS LENGTH); *F*(2.94, 55.94) = 1.579, *p* = .205, η_p_^2^ = .077.

## Discussion

In Experiment 1, we found that a single, innocuous electrocutaneous stimulus delivered to the left index finger inhibited CSE recorded from the right FDI-muscle at both early (i.e. 25 and 35 ms) and late (160 ms) intervals between tactile stimulation and M1-TMS. This finding is consistent with previous evidence of motor inhibition at short (Bailey et al., 2016; Bikmullina et al., 2009; Cash et al., 2015; Tamburin et al., 2001, 2005; Tamè et al., 2015; Tokimura et al., 2000; Udupa et al., 2009, 2014) and long (Chen et al., 1999; Sailer et al., 2002; Turco et al., 2017) interstimulus intervals. Furthermore, the observed recovery of motor excitability at a 45 ms interval suggests that the inhibitory effects of touch on motor functioning are time- locked and transient. We also found a facilitation effect at a 60 ms interstimulus interval, aligning with the results of Tamburin and colleagues (2001), who reported facilitated MEPs following electrocutaneous stimulation of the index finger at three times the sensory threshold (see also,Roy & Gorassini, 2008). Contrary to our hypothesis, the magnitude of afferent inhibition was not affected by the length of the afferent stimulus either at 0.2 or 0.4 ms. This suggests that the modulation of SAI by increased afferent recruitment is selective.

### Experiment 2: Corticospinal excitability and afferent inhibition as a function of tonic heat pain

Results of Experiment 1 demonstrated the validity of our approach and indicated that the stimulus length of the afferent signal did not affect SAI nor overall CSE. Given the effectiveness of this approach, we used a similar design in Experiment 2 to investigate the influence of tonic heat pain on the CSE. While previous studies suggest that thermal pain can modulate CSE (Delahunty et al., 2024; Martel et al., 2017; Mercier et al., 2016), evidence of such modulation in ipsilateral upper limb muscles is primarily limited to hand pain, with less consistent findings for forearm pain using saline, capsaicin and CO2-induced pain models (Larsen et al., 2018; Martel et al., 2017; Valeriani et al., 2001). Therefore, we tested whether cutaneous thermal pain applied to the forearm would significantly alter CSE, and whether these effects would occur independently of, or directly mediate, ongoing tactile sensorimotor interactions (i.e., SAI).

### Materials and Methods

#### Participants

Twenty new participants were recruited *(Mean_Age_*= 20.5 *SD_Age_*= 5.69 years; 6 males, 14 females). 15 were right-handed, 3 left-handed and 2 ambidextrous as recorded by the Edinburgh Handedness questionnaire (EHI; Oldfield, 1971; *M* = 55, *range* = -100-100). Recruitment methods and exclusion criteria were identical to Experiment 1.

### Thermal stimulation

Cutaneous tonic heat pain was induced on the participants’ left forearm by a 13 mm diameter contact thermal probe (NTE-2A Thermal Probe and controller; Physitemp Instruments LLC, Clifton NJ, USA) along with topical application of a 0.2% capsaicin cream. The topical application of low concentration capsaicin cream allows for sensitisation of C-nociceptive fibres for mild heat (<40°C) to be perceived as painful while minimizing the risk of tissue damage (Baumann et al., 1991). The thermode was integrated into the wooden desk with the probe tip flush with the tabletop (Figure 4A). This allowed for thermal stimulation to be delivered without additional tactile input elsewhere on the body

**Figure 4.**
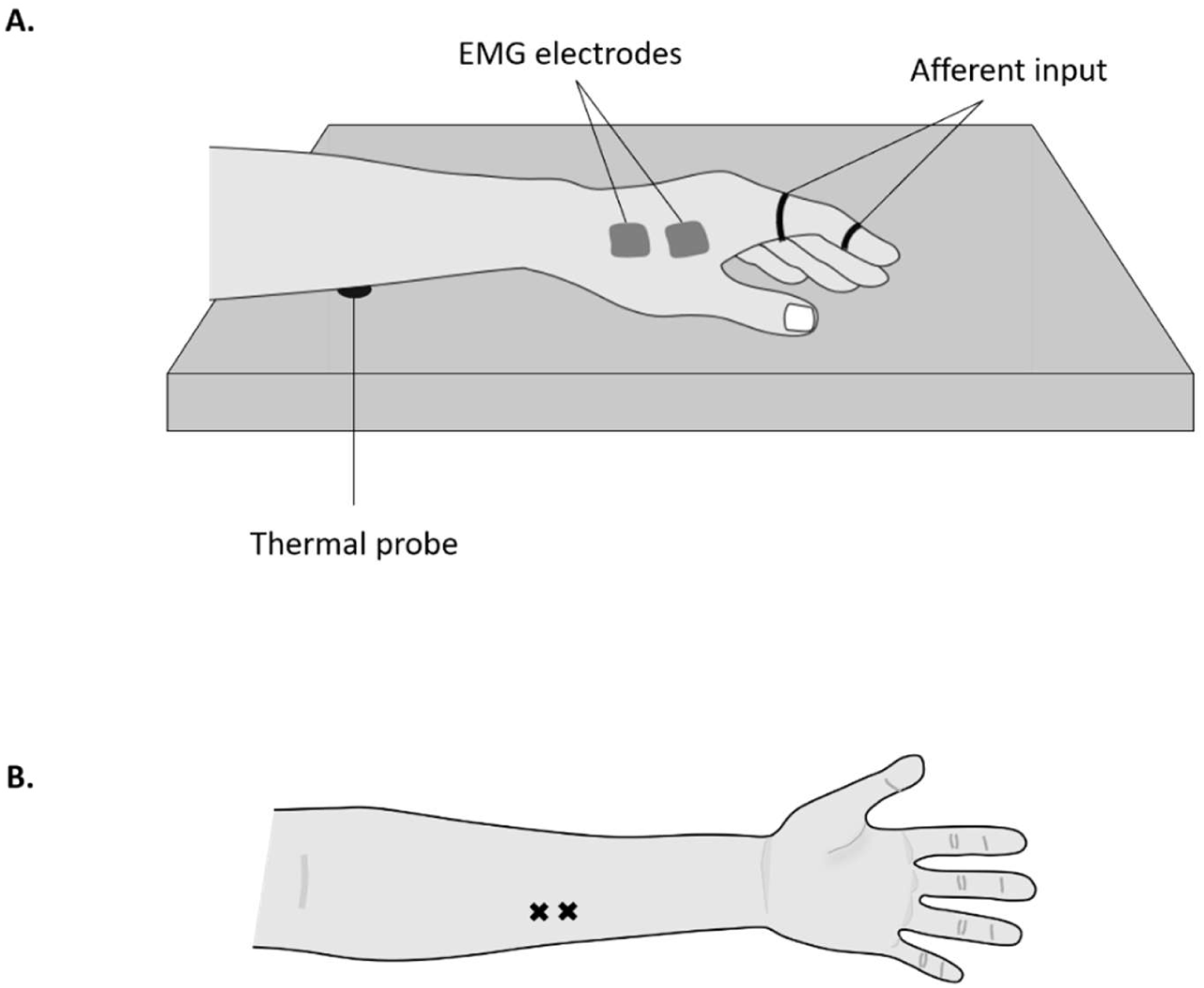
The experimental set-up. (A) Two points of thermode contact were marked on the right forearm. (B) Illustration of the set-up of thermal heat pain induction; the thermode was integrated into the desk surface and the arm remained rested on the probe throughout.

Two thermode stimulation points were marked on the participants arm, one midway between the centre-right of the cubital fossa and the wrist joint, and the second 2 cm distal to the first point (Figure 4B). These markers served as locations of thermal stimulation to avoid potential skin damage from prolonged thermode contact and, to minimize habituation or sensitisation over time. A half-pea-sized amount of capsaicin was applied to each marker for 10 minutes, then removed with a dry cloth before the start of the main experimental protocol. Five participants reported a mild but not painful warm and/or tingling sensation that resolved after the cream was removed. All other participants reported no perceptible sensations. Painful heat stimulation for the experimental protocol was calibrated to elicit a verbal rating of 7 on a pain scale (0 = “no pain at all”, 10 = “worst pain imaginable in the current context”). Starting at the baseline temperature (35°C), the temperature was manually increased in 0.5°C increments until the desired heat rating was attained.

### Experiment Protocol

The experimental protocol was identical to Experiment 1 with the following exception, continuous moderate heat pain was applied to the forearm throughout every block of trials. After TMS and electrotactile thresholding was performed, a moderate level of heat pain was calibrated. The thermode was moved between one of the two stimulation sites under an ABAB counterbalance design (Figure 4B), and the thermode temperature was adjusted in increments of 0.5^0^C to maintain a moderate level of pain (7/10). The mean average temperature delivered was 40.6°C (*SD = 3.6* °C). Thermode temperature required to elicit moderate pain significantly increased across blocks as indicated by a non-parametric Friedman test; *X^2^* (7) = 27.08, *p* = .001.

### Design

Given that Experiment 1 showed afferent durations of 0.2 and 0.4 ms had no effect on afferent inhibition, the electrotactile pulse width was arbitrarily set at 0.2 ms in accordance with previous reports (Chen et al., 1999; Sailer et al., 2002). The control group for comparison was taken from Experiment 1, Session 2. The effect of pain on SAI was examined on the within- participants factor of delay (DELAY: 15, 25, 35, 45 and 60 ms), and on the between-participants factor of pain condition (PAIN STATE: painful vs. painless). The effect of pain on CSE was assessed as a function of pain condition and on all conditions of afferent stimulation (CONDITION: TMS-only, 15, 25, 35, 45, 60 and 160 ms delays).

## Results

MEP ratios were calculated and log transformed as in Experiment 1. First, we compared differences in MEP ratios of the painful condition to a MEP ratio of 0 (TMS-only). Two-tailed t- tests showed that MEP ratios were significantly less than 0 at delays of 25 ms (*M* = -0.15, *SE* = 0.03; *t*(19) = 6.23, *p* = <.001, *d* = 1.39), 35 ms (*M* = -0.13, *SE* = 0.03; *t*(19) = 3.99, *p* = <.001, *d* = 0.89) and 160 ms (*M* = -0.28, *SE* = 0.05; *t*(19) = 5.22, *p* = <.001, *d* = 1.17). In contrast, at a 60 ms delay MEP ratios were significantly greater than 0 (*M* = 0.10, *SE* = 0.03; *t*(19) = -3.49, *p* = .002, *d* = -0.78.

Compound symmetry was significantly violated for the factor DELAY (Mauchly’s test: *p*<.001) and a Greenhouse Geiser correction was applied. A 2-way RM ANOVA was conducted on the factors DELAY and PAIN STATE, revealing a main effect of DELAY, *F*(2.85, 108.33) = 31.24, *p* <.001, η_p_^2^ = .451. However, there was no significant main effect of PAIN STATE, *F*(1,38) = 0.132, *p* = .719, η_p_^2^ = .003, and the two factors did not interact (DELAY X PAIN STATE), *F*(4, 152) = 0.54, *p* = .706, η_p_^2^ = .014 (Figure 5A). The effect of pain on LAI was assessed by an independent samples two-tailed t-test comparing MEP ratios at a 160ms delay between painless and painful conditions. This revealed no significant difference between the painful (*M* = -0.28, *SE* = 0.05) and painless (*M* = -0.35, *SE* = 0.05) conditions; *t*(38) = 0.92, *p* = .365, *d* = 0.3.

**Figure 5.**
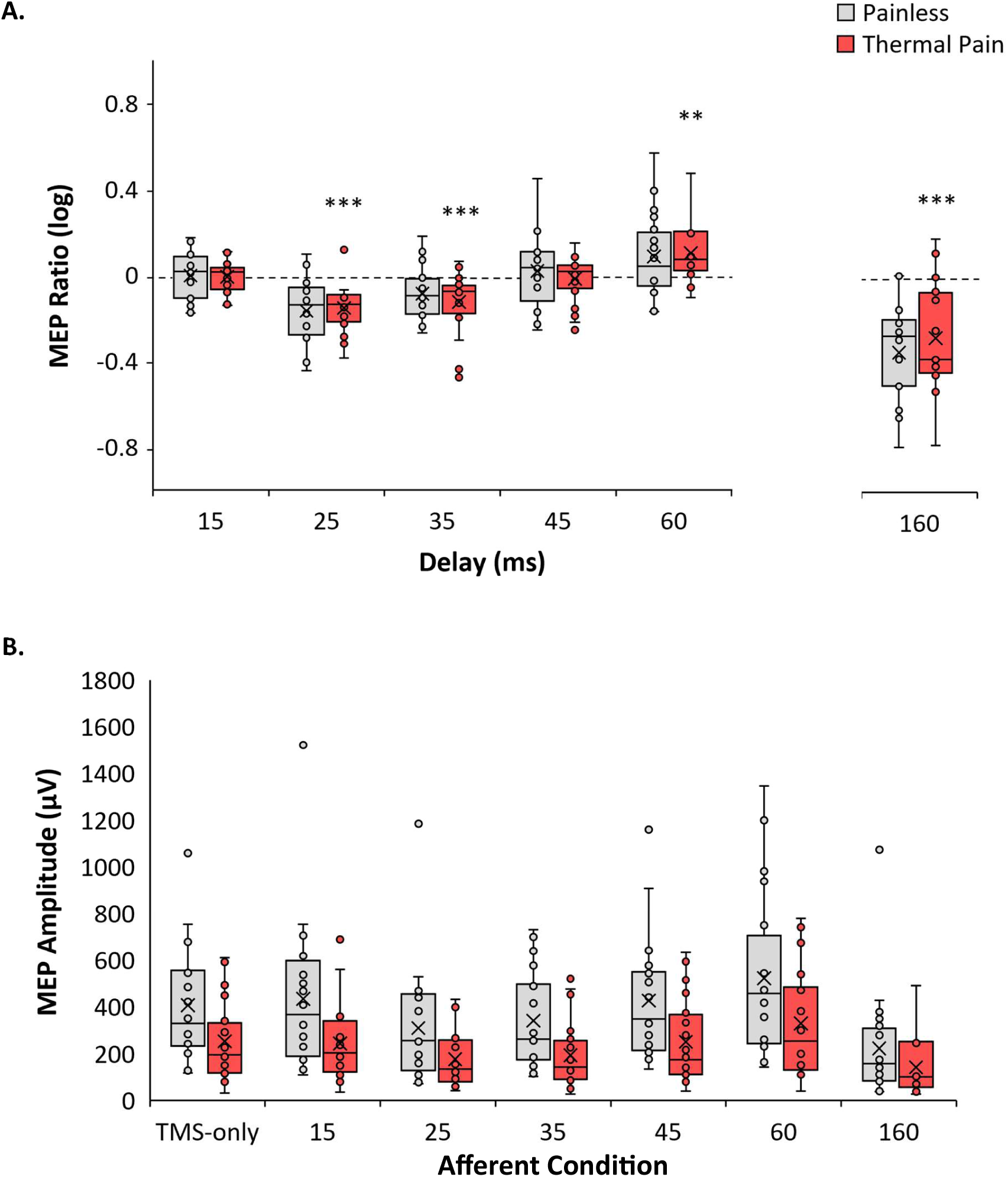
Boxplots show group data, centre crosses show the mean average and horizontal lines show exclusive median. Upper and lower whiskers show values within 1.5 times the interquartile range from lower and upper quartiles. Inner circle outlines show data points lying between inner and upper whiskers. (A) Showing MEP Ratios as a function of pain states and the delay (ms) between tactile input and M1-spTMS. Showing MEP Ratios as a function of pain states and the delay (ms) between tactile input and M1-spTMS. TMS-only trials correspond to a MEP ratio of 0. Boxplots show group data, centre crosses show the mean average and horizontal lines show exclusive median. Inner circle outlines show data points lying between inner and upper whiskers. \**P* < .05, \*\**P* < .01, \*\*\**P* < .001 indicate delays significantly different from a MEP ratio of 0 in the pain condition. (B) Raw MEP amplitudes across all trial types of afferent stimulation and TMS-only trials as a function of pain state.

The effect of pain on CSE was examined using a 2-way ANOVA on the factors of PAIN STATE and CONDITION, with a Greenhouse Geiser correction applied to the factor CONDITION (Mauchly’s test: *p* <.001). This revealed a significant main effect of CONDITION, *F*(2.87, 108.85) = 24.57, *p* <.001, η_p_^2^ = .393, reflecting the effects of inhibition and facilitation in MEP amplitudes before ratio corrections and transformations were applied. Whilst this did not vary as a function of pain (CONDITION x PAIN STATE, *F*(6, 228) = 1.50, *p* = .180, η_p_^2^ = .038), we found a significant main effect of PAIN STATE, *F*(1,38) = 5.78, *p* = .021, η_p_^2^ = .132. As shown in Figure 5B, MEP amplitudes were significantly lower in the painful (*M* = 226.64, *SE* = 45.34) compared to the painless group (*M* = 380.81, *SE* = 45.34; *t*(38) = -2.40, *p* = .021).

As a control check we compared tactile detection thresholds and TMS M1-FDI motor thresholds between the painful and painless conditions. A two-tailed t-tests revealed no significant difference in tactile detection thresholds (mA) between the painful (*M* = 1.42, *SE* = 0.1) and painless group (*M* = 1.53, *SE* = 0.07; *t*(38) = -0.88, *p* = .385, *d* = -0.28) and, no significant difference in motor thresholds (%) between the painful (*M* = 43.8, *SE* = 1.59) and painless (*M* = 42.4, *SE* =0.98; *t*(38) = 0.77, *p* = .449, *d* = 0.24) group.

## Discussion

In Experiment 2, we found that although moderate tonic heat pain to the ipsilateral forearm did not mediate the effects of tactile signalling on CSE, it did inhibit CSE across all conditions of tactile stimulation including TMS-only trials. This was indicated by significantly reduced MEP amplitudes in the painful compared to the painless group, demonstrating that while pain directly affects motor functioning, this relationship is not mediated through touch.

This finding demonstrate that pain does not influence either the temporal dynamics nor the magnitude of short- and long-afferent inhibition, as the results are comparable to what we found in painless condition. Our findings are consistent with that of Burns and colleagues (2016b) who reported no effect of saline-induced muscle pain in the tested muscle on SAI (Burns et al., 2016b). However, the present study extends this finding to cutaneous thermal pain applied to the forearm proximal to both the tactile input and tested muscle. Additionally, we broaden the scope of this finding across temporal delays. While Burns and colleagues (2016b) examined only 20 ms (SAI) and 200 ms (LAI) inter-stimulus intervals, we demonstrated that the preservation of afferent inhibition during tonic pain also applies to the afferent facilitation observed at a 60 ms delay and, to motor recovery at 45 ms.

## General Discussion

In the present study we used an AI paradigm to investigate the effects of tonic pain on corticospinal excitability and (tactile) sensorimotor interactions. We found that cutaneous heat pain applied to the forearm significantly reduced MEP amplitudes of the ipsilateral FDI muscle, signifying supressed motor output by pain. Despite the overall reduction in CSE, neither SAI or LAI were affected by pain. This aligns with some previous studies (Burns et al., 2016b; Mercier et al., 2016), though it contrasts with others such as Delahunty and colleagues (2024), who reported reduced SAI during painful cold water immersion of the whole contralateral hand (Delahunty et al., 2024). This discrepancy likely stems from considerable differences in the nature of painful stimulation. Additionally, we found no effect of pain on afferent facilitation (60 ms interstimulus interval) or MEP recovery (45 ms), thereby extending previous literature on the influence of pain on afferent inhibition. Collectively, our results indicate the whilst tonic pain can supress CSE, other sensorimotor interactions remain undisturbed. Therefore, our data supports a direct route by which pain influences motor response as shown in Figure 1A.

The finding that tonic heat pain exerts an inhibitory influence on motor output aligns with previous studies demonstrating reduced CSE in response to painful heat exposure (Billot et al., 2018; Cheong et al., 2003; Dubé & Mercier, 2011; Farina et al., 2001; Mavromatis et al., 2016; Mercier et al., 2016). Notably, we emphasise this finding in light of a recent meta- analysis suggesting limited evidence to show the effect of tonic forearm pain on CSE (Rohel et al., 2021). One possible explanation for the suppression of CSE is that the downregulation of corticomotor activity facilitates the prioritisation of adaptive spinal reflex responses (Farina et al., 2003; Inghilleri et al., 1997). Additionally, supressed CSE may reflect central nervous system mechanisms aimed at minimizing further pain and tissue damage (Hodges & Tucker, 2011).

Alternatively, we propose a role for executive control in maintaining limb and body posture against an ongoing noxious stimulus. In the current experiment, thermal stimulation was calibrated to elicit moderate pain (rated seven out of ten on the pain scale), and participants were instructed to remove the limb only if pain became intolerable - we note that this situation never occurred. The observed suppression of MEPs in the ipsilateral FDI may reflect the exertion of such top-down control in experimental contexts. This notion is supported by evidence showing that reductions in CSE are associated with lower pain severity (Chowdhury et al., 2022). Indeed, supressed CSE may subserve higher-order strategies in response to sustained painful input (see, for example, Legrain et al., 2011).

Across two experiments, this study replicates previous findings that TMS-evoked MEPs in the FDI muscle can be inhibited by a preceding electrical stimulus to the index finger of the ipsilateral hand (Bikmullina et al., 2009; Helmich et al., 2005; Tamburin et al., 2005; Tamè et al., 2015; Tokimura et al., 2000). More specifically, we found that early intervals (i.e. 25 and 35 ms) are associated with inhibition, followed by recovery at 45 ms, facilitation at 60 ms, and returning inhibition at 160 ms. The time-dependant modulation of CSE likely reflects processes of sensorimotor interaction that facilitate hand-object interactions (Johansson & Flanagan, 2009). Additionally, we showed that this effect is consistent regardless of whether tactile stimulus duration is 0.2 and 0.4 ms.

The facilitating effect we found at 60 ms, while consistent with some reports (Devanne et al., 2009; Roy & Gorassini, 2008; Tamburin et al., 2001), contradicts others (Devanne et al., 2009; Tamè et al., 2015), such as a study by Tamè and colleagues (2015) who found MEP inhibition of the FDI muscle at this interval. Although MEP facilitation has been shown to increase with stronger afferent input (Fischer & Orth, 2011), both the present study and Tamè et al delivered tactile stimulation at the same intensity namely, 2.5 times the sensory threshold. The discrepancy in the results might stem from differences in the location of the afferent stimulation relative to the target muscle. In Tamè et al. (2015), inhibition was found at a 60 ms delay when applied to the ipsilateral third digit, whereas, in our study, it was delivered to the second digit - the one that the function of the FDI is predominantly related to (Masquelet et al., 1986). Furthermore, we propose that the observed facilitation may be linked to the long latency reflex (LLR) pathway, which follows the H-reflex response 50-100 ms following limb displacement or submaximal nerve stimulation (Pruszynski & Scott, 2012). Consequently, the dynamics of afferent inhibition and facilitation depend not only on the nature of afferent input (e.g. amplitude), but also on its spatial and functional relationship to the relevant muscle.

When interpreting the findings of the current study it is important to note that MEP amplitudes alone cannot distinguish between spinal and supraspinal processes contributing to MEP amplitudes. While LAI is suggested to be cortically driven, as its timing exceeds that of spinal reflex responses (Chen et al., 1999), the origins of SAI remain less certain (Turco et al., 2018). However, several early studies that used H-reflex responses and transcranial electrical stimulation provided evidence that pain-related inhibition of CSE is primarily cortical (Farina et al., 2001; Valeriani et al., 1999, 2001), or results from a combination of cortical inhibition and later H-reflex modulation (Le Pera et al., 2001). This underscores the need for further research to clarify the interaction between spinal and supraspinal processes in shaping functional motor responses to both noxious and innocuous somatosensory inputs. In the present study the effect of thermal pain on CSE, SAI and LAI were assessed during painful stimulation, previous studies have shown that SAI and LAI can change following the cessation of pain (Burns et al., 2016b; Delahunty et al., 2024). Therefore, future studies should assess these phenomena in a post-pain context as this can provide insight into potential plasticity- related changes in the sensorimotor system after exposure to and recovery from noxious input.

## Conclusion

The present study demonstrated that moderate tonic heat pain applied to the left forearm reduced TMS-evoked MEP amplitudes in the left FDI muscle, while tactile afferent inhibition remained unchanged. This suggests the existence of a direct pathway between cutaneous noxious input and cortically driven motor output, operating independently of tactile sensorimotor interactions. The observed pain-induced inhibition of CSE aligns with existing literature; however, we further show that cutaneous forearm pain can alter motor activity in the distal hand. These findings, observed in healthy participants, may have important clinical implications for understanding how pain can directly affect motor processes whilst other networks of sensorimotor interactions remain unaffected. Furthermore, our paradigm offers an approach for investigating disrupted motor control and sensorimotor disturbances in different forms of chronic pain.

## Credit author statement

Louisa Gwynne: Conceptualization, Methodology, Validation, Formal analysis, Investigation, Writing – original draft, Visualization; Luigi Tamè: Conceptualization, Methodology, Validation, Formal analysis, Resources, Writing, review & editing.

## Acknowledgments

We would like to thank Nicholas Paul Holmes for his insightful comments on an early version of the manuscript. Katherine Dyke and Mimma Veniero for their insightful comments on the data. LG was supported by the South East Network for Social Sciences (SeNSS) Economic and Social Research Council (ESRC)-funded Doctoral Training Partnership.

## References

Allison, T., McCarthy, G., & Wood, C. C. (1992). The relationship between human long-latency somatosensory evoked potentials recorded from the cortical surface and from the scalp. Electroencephalography and Clinical Neurophysiology, 84(4), 301–314. 10.1016/0168-5597(92)90082-m

Bailey, A. Z., Asmussen, M. J., & Nelson, A. J. (2016). Short-latency afferent inhibition determined by the sensory afferent volley. Journal of Neurophysiology, 116(2), 637–644. 10.1152/jn.00276.2016

Baumann, T. K., Simone, D. A., Shain, C. N., & LaMotte, R. H. (1991). Neurogenic hyperalgesia: the search for the primary cutaneous afferent fibers that contribute to capsaicin-induced pain and hyperalgesia. Journal of Neurophysiology, 66(1), 212–227. 10.1152/jn.1991.66.1.212

Bikmullina, R., Kičić, D., Carlson, S., & Nikulin, V. V. (2009). Electrophysiological correlates of short-latency afferent inhibition: A combined EEG and TMS study. Experimental Brain Research, 194(4), 517–526. 10.1007/s00221-009-1723-7

Billot, M., Neige, C., Gagné, M., Mercier, C., & Bouyer, L. J. (2018). Effect of Cutaneous Heat Pain on Corticospinal Excitability of the Tibialis Anterior at Rest and during Submaximal Contraction. Neural Plasticity, 2018. 10.1155/2018/8713218

Burns, E., Chipchase, L. S., & Schabrun, S. M. (2016a). Primary sensory and motor cortex function in response to acute muscle pain: A systematic review and meta-analysis. European Journal of Pain (United Kingdom), 20(8), 1203–1213. 10.1002/ejp.859

Burns, E., Chipchase, L. S., & Schabrun, S. M. (2016b). Reduced short- and long-latency afferent inhibition following acute muscle pain: A potential role in the recovery of motor output. Pain Medicine (United States), 17(7), 1343–1352. 10.1093/pm/pnv104

Cash, R. F. H., Isayama, R., Gunraj, C. A., Ni, Z., & Chen, R. (2015). The influence of sensory afferent input on local motor cortical excitatory circuitry in humans. Journal of Physiology, 593(7), 1667–1684. 10.1113/jphysiol.2014.286245

Chen, R., Corwell, B., & Hallett, M. (1999). Modulation of motor cortex excitability by median nerve and digit stimulation. Experimental Brain Research, 129(1), 77–86. 10.1007/s002210050938

Cheong, J. Y., Yoon, T. S., & Lee, S. J. (2003). Evaluations of inhibitory effect on the motor cortex by cutaneous pain via application of capsaicin. Electromyography and Clinical Neurophysiology, 43(4), 203–210.

Chipchase, L., Schabrun, S., Cohen, L., Hodges, P., Ridding, M., Rothwell, J., Taylor, J., & Ziemann, U. (2012). A checklist for assessing the methodological quality of studies using transcranial magnetic stimulation to study the motor system: An international consensus study. Clinical Neurophysiology, 123(9), 1698–1704. 10.1016/j.clinph.2012.05.003

Chipchase, L., Schabrun, S., & Hodges, P. (2011). Peripheral electrical stimulation to induce cortical plasticity: A systematic review of stimulus parameters. Clinical Neurophysiology, 122(3), 456–463. 10.1016/j.clinph.2010.07.025

Choudhury, S., Siddique, U., Rahman, S., Kumar, Y., Banerjee, S., Baker, M. R., Baker, S. N., & Kumar, H. (2023). Short-Latency Afferent Inhibition Correlates with Stage of Disease in Parkinson’s Patients. Canadian Journal of Neurological Sciences, 50(4), 579–583. 10.1017/cjn.2022.83

Chowdhury, N. S., Chang, W. J., Millard, S. K., Skippen, P., Bilska, K., Seminowicz, D. A., & Schabrun, S. M. (2022). The Effect of Acute and Sustained Pain on Corticomotor Excitability: A Systematic Review and Meta-Analysis of Group and Individual Level Data. Journal of Pain, 23(10), 1680–1696. 10.1016/j.jpain.2022.04.012

Delahunty, E. T., Bisset, L. M., & Kavanagh, J. J. (2024). Short-latency afferent inhibition is reduced with cold-water immersion of a limb and remains reduced after removal from the cold stimulus. Experimental Physiology, March, 1817–1825. 10.1113/EP091896

Devanne, H., Degardin, A., Tyvaert, L., Bocquillon, P., Houdayer, E., Manceaux, A., Derambure, P., & Cassim, F. (2009). Afferent-induced facilitation of primary motor cortex excitability in the region controlling hand muscles in humans. European Journal of Neuroscience, 30(3), 439–448. 10.1111/j.1460-9568.2009.06815.x

Di Lazzaro, V., Oliviero, A., Tonali, P. A., Marra, C., Daniele, A., Profice, P., Saturno, E., Pilato, F., Masullo, C., & Rothwell, J. C. (2002). Noninvasive in vivo assessment of cholinergic cortical circuits in AD using transcranial magnetic stimulation. Neurology, 59(3), 392–397. 10.1212/WNL.59.3.392

Dubé, J. A., & Mercier, C. (2011). Effect of pain and pain expectation on primary motor cortex excitability. Clinical Neurophysiology, 122(11), 2318–2323. 10.1016/j.clinph.2011.03.026

Farina, S., Tinazzi, M., Le Pera, D., & Valeriani, M. (2003). Pain-related modulation of the human motor cortex. Neurological Research, 25(2), 130–142. 10.1179/016164103101201283

Farina, S., Valeriani, M., Rosso, T., Aglioti, S., Tamburin, S., Fiaschi, A., & Tinazzi, M. (2001). Transient inhibition of the human motor cortex by capsaicin-induced pain. A study with transcranial magnetic stimulation. Neuroscience Letters, 314(1–2), 97–101. 10.1016/S0304-3940(01)02297-2

Faul, F., Erdfelder, E., Lang, A.-G., & Buchner, A. (2007). G*Power 3: A flexible statistical power analysis program for the social, behavioral, and biomedical sciences. Behavior Research Methods, 39(2), 175–191. 10.3758/BF03193146

Fischer, M., & Orth, M. (2011). Short-latency sensory afferent inhibition: Conditioning stimulus intensity, recording site, and effects of 1 Hz repetitive TMS. Brain Stimulation, 4(4), 202–209. 10.1016/j.brs.2010.10.005

Hamada, Y., Otsuka, S., Okamoto, T., & Suzuki, R. (2002). The profile of the recovery cycle in human primary and secondary somatosensory cortex: A magnetoencephalography study. Clinical Neurophysiology, 113(11), 1787–1793. 10.1016/S1388-2457(02)00258-4

Harris, A. J. (1999). Cortical origin of pathological pain. Lancet, 354(9188), 1464–1466. 10.1016/S0140-6736(99)05003-5

Helmich, R. C. G., Bäumer, T., Siebner, H. R., Bloem, B. R., & Münchau, A. (2005). Hemispheric asymmetry and somatotopy of afferent inhibition in healthy humans. Experimental Brain Research, 167(2), 211–219. 10.1007/s00221-005-0014-1

Hodges, P. W., & Tucker, K. (2011). Moving differently in pain: A new theory to explain the adaptation to pain. In Pain (Vol. 152, Issue SUPPL.3). 10.1016/j.pain.2010.10.020

Inghilleri, M., Cruccu, G., Argenta, M., Polidori, L., & Manfredi, M. (1997). Silent period in upper limb muscles after noxious cutaneous stimulation in man. In Electroencephalography and clinical Neurophysiology (Vol. 105).

Johansson, R. S., & Flanagan, J. R. (2009). Coding and use of tactile signals from the fingertips in object manipulation tasks. In Nature Reviews Neuroscience (Vol. 10, Issue 5, pp. 345–359). 10.1038/nrn2621

Larsen, D. B., Graven-Nielsen, T., Hirata, R. P., & Boudreau, S. A. (2018). Differential Corticomotor Excitability Responses to Hypertonic Saline-Induced Muscle Pain in Forearm and Hand Muscles. Neural Plasticity, 2018. 10.1155/2018/7589601

Larsen, D. B., Graven-Nielsen, T., Hirata, R. P., Seminowicz, D., Schabrun, S., & Boudreau, S. A. (2019). Corticomotor excitability reduction induced by experimental pain remains unaffected by performing a working memory task as compared to staying at rest. Experimental Brain Research, 237(9), 2205–2215. 10.1007/s00221-019-05587-y

Le Pera, D., Graven-Nielsen, T., Valeriani, M., Oliviero, A., Lazzaro, V. Di, Tonali, P. A., & Arendt-Nielsen, L. (2001). Inhibition of motor system excitability at cortical and spinal level by tonic muscle pain. www.elsevier.com/locate/clinph

Legrain, V., Iannetti, G. D., Plaghki, L., & Mouraux, A. (2011). The pain matrix reloaded: A salience detection system for the body. Progress in Neurobiology, 93(1), 111–124. 10.1016/j.pneurobio.2010.10.005

Levitt, H. (1971). Transformed up-down methods in psychoacoustics. The Journal of the Acoustical Society of America, 49(2B), 467–477.

Martel, M., Harvey, M. P., Houde, F., Balg, F., Goffaux, P., & Léonard, G. (2017). Unravelling the effect of experimental pain on the corticomotor system using transcranial magnetic stimulation and electroencephalography. Experimental Brain Research, 235(4), 1223– 1231. 10.1007/s00221-017-4880-0

Masquelet, A. C., Salama, J., Outrequin, G., Serrault, M., & Chevrel, J. P. (1986). Morphology and functional anatomy of the first dorsal interosseous muscle of the hand. Surgical and Radiologic Anatomy : SRA, 8(1), 19–28. 10.1007/BF02539704

Mauguière, F., Merlet, I., Forss, N., Vanni, S., Jousmäki, V., Adeleine, P., & Hari, R. (1997). Activation of a distributed somatosensory cortical network in the human brain: a dipole modelling study of magnetic fields evoked by median nerve stimulation. Part II: Effects of stimulus rate, attention and stimulus detection. Electroencephalography and Clinical Neurophysiology, 104(4), 290–295. 10.1016/s0013-4694(97)00018-7

Mavromatis, N., Gagné, M., Voisin, J. I. A. V., Reilly, K. T., & Mercier, C. (2016). Experimental tonic hand pain modulates the corticospinal plasticity induced by a subsequent hand deafferentation. Neuroscience, 330, 403–409. 10.1016/j.neuroscience.2016.06.008

McCabe, C. S., Haigh, R. C., Halligan, P. W., & Blake, D. R. (2005). Simulating sensory-motor incongruence in healthy volunteers: Implications for a cortical model of pain. Rheumatology, 44(4), 509–516. 10.1093/rheumatology/keh529

Mercier, C., Gagné, M., Reilly, K. T., & Bouyer, L. J. (2016). Effect of experimental cutaneous hand pain on corticospinal excitability and short afferent inhibition. Brain Sciences, 6(4). 10.3390/brainsci6040045

Oldfield, R. C. (1971). The assessment and analysis of handedness: the Edinburgh inventory. In Neuropsychologia (Vol. 9). Pergamon Press.

Pruszynski, J. A., & Scott, S. H. (2012). Optimal feedback control and the long-latency stretch response. In Experimental Brain Research (Vol. 218, Issue 3, pp. 341–359). 10.1007/s00221-012-3041-8

Rice, D. A., Lewis, G. N., Graven-Nielsen, T., Luther, R., & McNair, P. J. (2021). Experimental Hand and Knee Pain Cause Differential Effects on Corticomotor Excitability. Journal of Pain, 22(7), 789–796. 10.1016/j.jpain.2021.01.006

Rohel, A., Bouffard, J., Patricio, P., Mavromatis, N., Billot, M., Roy, J. S., Bouyer, L., Mercier, C., & Masse-Alarie, H. (2021). The effect of experimental pain on the excitability of the corticospinal tract in humans: A systematic review and meta-analysis. European Journal of Pain (United Kingdom), 25(6), 1209–1226. 10.1002/ejp.1746

Romaniello, A., Cruccu, G., McMillan, A. S., Arendt-Nielsen, L., & Svensson, P. (2000). Effect of experimental pain from trigeminal muscle and skin on motor cortex excitability in humans. Brain Research, 882(1–2), 120–127. 10.1016/S0006-8993(00)02856-0

Rossi, S., Antal, A., Bestmann, S., Bikson, M., Brewer, C., Brockmöller, J., Carpenter, L. L., Cincotta, M., Chen, R., Daskalakis, J. D., Di Lazzaro, V., Fox, M. D., George, M. S., Gilbert, D., Kimiskidis, V. K., Koch, G., Ilmoniemi, R. J., Pascal Lefaucheur, J., Leocani, L., … Hallett, M. (2021). Safety and recommendations for TMS use in healthy subjects and patient populations, with updates on training, ethical and regulatory issues: Expert Guidelines. Clinical Neurophysiology, 132(1), 269–306. 10.1016/j.clinph.2020.10.003

Roy, F. D., & Gorassini, M. A. (2008). Peripheral sensory activation of cortical circuits in the leg motor cortex of man. Journal of Physiology, 586(17), 4091–4105. 10.1113/jphysiol.2008.153726

Sailer, A., Molnar, G. F., Cunic, D. I., & Chen, R. (2002). Effects of peripheral sensory input on cortical inhibition in humans. Journal of Physiology, 544(2), 617–629. 10.1113/jphysiol.2002.028670

Sailer, A., Molnar, G. F., Paradiso, G., Gunraj, C. A., Lang, A. E., & Chen, R. (2003). Short and long latency afferent inhibition in Parkinson’s disease. Brain, 126(8), 1883–1894. 10.1093/brain/awg183

Schabrun, S. M., & Hodges, P. W. (2012). Muscle pain differentially modulates short interval intracortical inhibition and intracortical facilitation in primary motor cortex. Journal of Pain, 13(2), 187–194. 10.1016/j.jpain.2011.10.013

Tamburin, S., Fiaschi, A., Andreoli, A., Marani, S., & Zanette, G. (2005). Sensorimotor integration to cutaneous afferents in humans: the effect of the size of the receptive field. Experimental Brain Research. Experimentelle Hirnforschung. Expérimentation Cérébrale, 167(3), 362–369. 10.1007/s00221-005-0041-y

Tamburin, S., Manganotti, P., Zanette, G., & Fiaschi, A. (2001). Cutaneomotor integration in human hand motor areas: Somatotopic effect and interaction of afferents. Experimental Brain Research, 141(2), 232–241. 10.1007/s002210100859

Tamè, L., & Holmes, N. P. (2023). Neurostimulation in Tactile Perception (pp. 451–482). 10.1007/978-1-0716-3068-6_20

Tamè, L., Pavani, F., Braun, C., Salemme, R., Farnè, A., & Reilly, K. T. (2015). Somatotopy and temporal dynamics of sensorimotor interactions: Evidence from double afferent inhibition. European Journal of Neuroscience, 41(11), 1459–1465. 10.1111/ejn.12890

Tokimura, H., Lazzaro, V. Di, Tokimura, Y., Oliviero, A., Profice, P., Insola, A., Mazzone, P., Tonali, P., & Rothwell, J. C. (2000). Tokimura et al. (2000). Journal of Physiology, *523*(2), 503–513.

Turco, C. V., El-Sayes, J., Fassett, H. J., Chen, R., & Nelson, A. J. (2017). Modulation of long- latency afferent inhibition by the amplitude of sensory afferent volley. Journal of Neurophysiology, 118(1), 610–618. 10.1152/jn.00118.2017

Turco, C. V., El-Sayes, J., Savoie, M. J., Fassett, H. J., Locke, M. B., & Nelson, A. J. (2018). Short- and long-latency afferent inhibition; uses, mechanisms and influencing factors. In Brain Stimulation (Vol. 11, Issue 1, pp. 59–74). Elsevier Inc. 10.1016/j.brs.2017.09.009

Udupa, K., Ni, Z., Gunraj, C., & Chen, R. (2009). Interactions between short latency afferent inhibition and long interval intracortical inhibition. Experimental Brain Research, 199(2), 177–183. 10.1007/s00221-009-1997-9

Udupa, K., Ni, Z., Gunraj, C., & Chen, R. (2014). Downloaded from journals.physiology.org/journal/jn at Univ of Kent Templeman Lib. J Neuro-Physiol, 111, 1350–1361. 10.1152/jn.00613.2013.-Peripheral

Valeriani, M., Restuccia, D., Di Lazzaro, V., Oliviero, A., Le Pera, D., Profice, P., Saturno, E., & Tonali, P. (2001). Inhibition of biceps brachii muscle motor area by painful heat stimulation of the skin. Experimental Brain Research, 139(2), 168–172. 10.1007/s002210100753

Valeriani, M., Restuccia, D., Lazzaro, V. Di, Oliviero, A., Profice, P., Le Pera, D., Saturno, E., & Tonali, P. (1999). Inhibition of the human primary motor area by painful heat stimulation of the skin.

Widener, G. L., & Cheney, P. D. (1997). Effects on muscle activity from microstimuli applied to somatosensory and motor cortex during voluntary movement in the monkey. Journal of Neurophysiology, 77(5), 2446–2465. 10.1152/jn.1997.77.5.2446

Yarnall, A. J., Rochester, L., Baker, M. R., David, R., Khoo, T. K., Duncan, G. W., Galna, B., & Burn, D. J. (2013). Short latency afferent inhibition: A biomarker for mild cognitive impairment in Parkinson’s disease? Movement Disorders, 28(9), 1285–1288. 10.1002/mds.25360

Zafarana, A., Muret, D., Farnè, A., & Tamè, L. (2024). Somatosensory impact on motor cortex: how touch shapes motor behaviour.

